# Chemical proteomics reveals that the anticancer drug everolimus affects ubiquitin-proteasome system

**DOI:** 10.1101/2023.10.15.562182

**Authors:** Anna A. Lobas, Amir Ata Saei, Hezheng Lyu, Roman A. Zubarev, Mikhail V. Gorshkov

**Affiliations:** V.L. Talrose Institute for Energy Problems of Chemical Physics, Moscow, Russian Federation; Division of Physiological Chemistry I, Department of Medical Biochemistry and Biophysics, Karolinska Institutet, SE-17 177 Stockholm, Sweden; Biozentrum, University of Basel, 4056 Basel, Switzerland; Department of Microbiology, Tumor and Cell Biology, Karolinska Institutet, SE-17 177 Stockholm, Sweden; The National Medical Research Center for Endocrinology, Moscow, Russia

**Keywords:** drug treatment, cell line, expression proteomics, rapamycin, everolimus, ubiquitin-proteasome system, ubiquitination, diglycine modification, semi-tryptic peptides

## Abstract

Rapamycin is a natural antifungal, immunosuppressive, and antiproliferative compound allosterically inhibiting mTOR complex 1. The ubiquitin-proteasome system (UPS) responsible for protein turnover is usually not listed among the pathways affected by mTOR signaling; however, a number of reports indicated the interplay between UPS and mTOR. It is also reported that rapamycin and analogs can allosterically inhibit the proteasome itself. We studied the molecular effect of rapamycin and its analogs (rapalogs), everolimus, and temsirolimus by expression chemical proteomics on A549 cell line. The analysis revealed that the cellular response to everolimus differs dramatically from that of rapamycin and temsirolimus. In cluster analysis the effect of everolimus was similar to that of bortezomib, a well-established proteasome inhibitor. UPS-related pathways were enriched in the cluster of proteins specifically up-regulated upon everolimus and bortezomib treatments, suggesting that both compounds have similar proteasome inhibition effects. In particular, the total amount of ubiquitin was significantly elevated in the samples treated with everolimus and bortezomib, and the analysis of the polyubiquitination patterns revealed elevated intensities of the ubiquitin peptide with a GG modification at K48 residue, consistent with a bottleneck in the proteasomal protein degradation. Moreover, everolimus treatment resulted in both ubiquitin phosphorylation and a significant amount of semi-tryptic peptides illustrating the induction of protease activity. These observations suggest that everolimus affects the ubiquitin-proteasome system in a unique way and its mechanism of action is strikingly different from that of its close chemical analogs, rapamycin and temsirolimus. Data are available via ProteomeXchange with identifier PXD045774.

## I. Introduction

Hyperactivation of mammalian TOR (mTOR) complex 1 (mTORC1) is observed in numerous human cancers due to gain-of-function mutations in oncogenes, such as PI3K and AKT, and/or loss-of-function mutations in tumor suppressors.^1^ Rapamycin (sirolimus) is a natural compound known in cancer therapy for its antiproliferative properties acting as an allosteric inhibitor of mTORC1.^1^ The two rapamycin analogs (rapalogs) everolimus and temsirolimus, are believed to share the same mechanism of action^2^. Due to the numerous processes regulated by mTOR, its inhibition has multiple effects on the protein synthesis, cell cycle, autophagy, etc.^3^ Specifically, rapalogs inhibit cell proliferation and induce cell cycle arrest with accumulation of cells in G1 phase that is reportedly behind their anticancer efficacy.^2^

Different rapalogs have distinct pharmacokinetics and bioavailability. For example, the bioavailability of everolimus and temsirolimus is higher than that of rapamycin, but their half-life in blood is shorter (26–30 h and 9–27 h, vs. 46–78 h).^4,5^ These differences might explain the various clinical applications of these drugs.^6–8^

While rapamycin’s mechanism of action includes mainly inhibition of mTORC1^3^, it is speculated that its long-term administration leads to mTORC2 inhibition^1^, resulting in numerous side-effects, including the impact on glucose homeostasis. It was shown in a mice model that, presumably due to shorter blood half-life, the use of everolimus and temsirolimus instead of rapamycin may have significantly lower impact on glucose homeostasis.^9^ On the other hand, a cell-based study showed similar effect on mTORC2 complex formation by all three compounds^10^, and despite different pharmacokinetic and pharmacodynamic properties, rapamycin analogs showedin the clinica similar toxicity profiles to that of the parent compound.^11^

The ubiquitin-proteasome system (UPS) is the crucial pathway for intracellular protein degradationregulating homeostasis and various cellular events, including those involved in carcinogenesis.^12–17^ In eukaryotic cells, targeted proteins are marked by polyubiquitin followed by their degradation into peptides by the proteasome.^18,19^ Many of the proteins that undergo the ubiquitin-dependent proteolysis are the regulators of physiological and/or pathological processes in the cells and their degradation plays an essential role in the cell cycle progression, differentiation, proliferation, as well as apoptosis and mitosis.^14,20–22^ Furthermore, the UPS is responsible for the degradation of misfolded and mutated proteins^23^, thus ensuring normal cell functioning. On the one hand, malfunctioning UPS may cause aberrant regulation of cell cycle proteins, which may affect the cell division in an uncontrolled manner, further leading to carcinogenesis.^24–26^ On the other hand, proteasome inhibition may exhibit potential as an anticancer treatment by targeting the protein function responsible for tumor growth and progression.^15,27,28^ In particular, proteasome inhibitors confine cancer progression by interfering with the temporal degradation of the regulatory proteins, thus sensitizing cancer cellsto apoptosis.^29^ While targeting the proteasome seems counterintuitive at first glance because of the essential role of the UPS in cellular homeostasis, a number of studies demonstrated that proteasome inhibition may lead to the accumulation of pro-apoptotic proteins in cancer rather than in normal cells.^30–32^ Indeed, some earlier reports indicated possible cytotoxic effects of proteasome inhibitors^33,34^, and searching for the ones with potential for cancer treatment has been the subject of intense investigations in drug discovery^35,36^, including the recent studies on drug repurposing.^37–41^

In the rather short list of proteasome inhibitors with clinical potential for cancer treatment, bortezomib was the first one approved by the US FDA for the treatment of multiple myeloma and mantle cell lymphoma^42–45^ One of the proposed mechanisms of its antitumor activity is the promotion of degradation of anti-apoptotic proteins and prohibiting the degradation of pro-apoptotic proteins through the proteasome inhibition that together result in the tumor cell death. The approval for clinical use of proteasome inhibitors confirmed that targeting the UPS is a feasible approach to treating different types of cancers.^46–48^ However, in addition to the toxic side effects, the use of currently approved proteasome inhibitors in cancer treatment is limited due to drug resistance, for example tobortezomib.^49,50^ This stimulated the on-going efforts to identify and/or develop the next generation of anti-cancer drugs targeting the UPS.^51–54^

Since both mTOR and UPS play a key role in protein turnover, there are continued debates on the possible impact on the proteasome function by mTOR inhibitors, such as rapamycin and rapalogs. It remains unclear, for example, whether the UPS-based proteolysis increases when mTOR activity decreases.^55,56^ Different reports provide contradictory views on the issue, one suggesting that mTORC1 inhibition reduces proteolysis by suppressing proteasome expression, while the others posit that mTOR inhibition results in increasing the cellular content of K48-linked ubiquitinated proteins and, thus, enhances the UPS-dependent proteolysis.^55^ There is also a study suggesting that rapamycin can allosterically inhibit the proteasome itself.^56^

In order to better understand the mechanistic similarities and differences of the molecular effects of rapamycin and its analogs (rapalogs), in this work we performed expression proteomic analysis of A549 cell line treated with 4 drugs, including rapamycin and its analogs (rapalogs), everolimus, and temsirolimus, as well as bortezomib, followed by multiplexed quantitative proteome analyses.^57^ The study was further extended by re-analysis of publicly available expandable proteome signature library of 56 anticancer molecules in cancer cell lines, ProTargetMiner^58^, with specific purpose of searching for the drugs exhibiting the proteasome inhibitor activity. Here we show that unlike rapamycin and temsirolimus, everolimus affects the UPS system.

## II. Materials and Methods

### Sample preparation

Human A549 cells (ATCC, USA) were grown in DMEM medium (Fisher Scientific) supplemented with 10% FBS (Fisher Scientific), 2 mM L-glutamine (Fisher Scientific) and 100 units per mL of penicillin/streptomycin (Thermo Fisher) and incubated at 37 °C in 5% CO_2_. Rapamycin (S1039), everolimus (S1120), temsirolimus (S1044) and bortezomib (S1013) were purchased from Selleckchem as 10 mM solutions in DMSO.

In LC50 determination, cells were seeded at a density of 4,000 per well in 96-well plates and after a day of growth, were treated with the serial concentrations of molecules for 48 h. Thereafter cell viability was measured using CellTiter-Blue Cell Viability Assay (Promega; Cat#G8081) according to the manufacturer protocol.

Cells were seeded at a density of 250,000 per well and allowed to grow for 24 h in biological triplicates. Next, cells were either treated with vehicle (DMSO) or compounds at IC50 concentrations (25 μM for rapamycin and temsirolimus, 50 μM for everolimus, and 0.15 μM for bortezomib) for 48h. After treatment, cells were collected, washed twice with PBS (Fisher Scientific) and then lysed using 8 M urea, 1% SDS, and 50 mM Tris at pH 8.5 with protease inhibitors (Sigma; Cat#05892791001). The cell lysates were subjected to 1 min sonication on ice using Branson probe sonicator and 3 s on/off pulses with a 30% amplitude. Protein concentration was then measured for each sample using a BCA Protein Assay Kit (Thermo) and 25 µg of each sample was reduced with DTT (final concentration 10 mM) (Sigma; Cat#D0632) for 1 h at room temperature. Afterwards, iodoacetamide (IAA) (Sigma; Cat#I6125) was added to a final concentration of 50 mM. The samples were incubated at room temperature for 1 h in the dark, with the reaction being stopped by addition of 10 mM DTT. After precipitation of proteins using methanol/chloroform, the semi-dry protein pellet was dissolved in 25 µL of 8 M urea in 20 mM EPPS (pH 8.5) (Sigma; Cat#E9502) and was then diluted with EPPS buffer to reduce urea concentration to 4 M. Lysyl endopeptidase (LysC) (Wako; Cat#125-05061) was added at a 1: 75 w/w ratio to protein and incubated at room temperature overnight. After diluting urea to 1 M, trypsin (Promega; Cat#V5111) was added at the ratio of 1: 75 w/w and the samples were incubated for 6 h at room temperature.

Acetonitrile (Fisher Scientific; Cat#1079-9704) was added to a final concentration of 20% v/v. TMTpro16 reagents (Thermo; Cat#90110) were added 4x by weight to each sample, followed by incubation for 2 h at room temperature. The reaction was quenched by addition of 0.5% hydroxylamine (Thermo Fisher; Cat#90115). Samples were combined, acidified by trifluoroacetic acid (TFA; Sigma; Cat#302031-M), cleaned using Sep-Pak (Waters; Cat#WAT054960) and dried using a DNA 120 SpeedVac™ concentrator (Thermo).

The pooled dried sample was resuspended in 20 mM ammonium hydroxide and separated into 96 fractions on an XBrigde BEH C18 2.1 × 150 mm column (Waters; Cat#186003023), using a Dionex Ultimate 3000 2DLC system (Thermo Scientific) over a 48 min gradient of 1–63%B (*B* = 20 mM ammonium hydroxide in acetonitrile) in three steps (1–23.5%B in 42 min, 23.5–54%B in 4 min and then 54–63%B in 2 min) at 200 µL min^−1^ flow. Fractions were then concatenated into 24 fractions (e.g. A1, C1, E1, G1). After drying and resuspension in 0.1% formic acid (FA) (Fisher Scientific), each fraction was analyzed over a 100 min gradient (total method time = 120 min) in random order.

### Chemical proteomics analysis

Samples were loaded with buffer A (0.1% FA in water) onto a 50 cm EASY-Spray column (75 µm internal diameter, packed with PepMap C18, 2 µm beads, 100 Å pore size) connected to a nanoflow Dionex UltiMate 3000 UPLC system (Thermo) and eluted in an increasing organic solvent gradient from 4 to 28% over 90 min and up to 34% till 100 min (B: 98% ACN, 0.1% FA, 2% H2O) at a flow rate of 300 nL min−1. Mass spectra were acquired with an Orbitrap Lumos mass spectrometer (Thermo) in data-dependent mode with MS1 scan at 120,000 resolution, and MS2 at 50,000 (@200 m/z), in the mass range from 400-1600 m/z. Isolation window was set at 1.6. Peptide fragmentation was performed via higher-energy collision dissociation (HCD) with energy set at 35 NCE. The mass spectrometry proteomics data have been deposited to the ProteomeXchange Consortium via the PRIDE ^59^ partner repository with the dataset identifier PXD045774.

### Data source

Additional data analysis was also performed on a previously published ProTargetMiner dataset.^58^ This dataset of 229 LC-MS/MS analyses presents the results of expression proteomics screening on A549 cells treated with 56 anticancer agents.

### Data analysis

The raw data from TMT-based LC-MS/MS were converted to mzML by MSconvert and analyzed by IdentiPy, version 0.3.7^60^ with a post-search validation performed using Scavager, version 0.2.12^61^. MS/MS data were searched against the SwissProt protein sequence database (human, version 04_2021, 20395 entries). Cysteine carbamidomethylation was used as a fixed modification, while methionine oxidation and protein N-terminal acetylation were selected as variable modifications. Trypsin was selected as enzyme specificity. No more than two missed cleavages were allowed. A 1% false discovery rate was used as a filter at both protein and peptide levels. The molecular mass tolerance was 10 ppm, and the minimum peptide length was 5 residues. For the analysis of proteolysis products, a semi-tryptic search with similar parameters was performed. For the polyubiquitination search, a limited database containing only ubiquitin sequences with different sites of ubiquitination (addition of GG residues to K residue, Supplementary Table 1) was used with the same search settings with an addition of serine phosphorylation potential modification. Post-search validation, as well as extraction and quantification of the TMT reporter ions in the mass spectra were performed using Scavager. Quantitation results were normalized by the sum of all intensities within a TMT channel. Gene ontology (GO) enrichment for biological processes was performed using Enrichr^62^ (https://maayanlab.cloud/Enrichr/), the set of genes corresponding to all identified proteins was used as the background. Gene ontology (GO) terms with adjusted p-values < 0.05 were considered. Motif enrichment was performed using an in-house script comparing the amino acid frequencies at each position to that of random background samples with a p-value threshold of 0.05.

## III. Results and discussion

### Overview of A549 proteome upon treatment

We performed multiplex expression proteomics analysis of A549 cells treated by rapamycin, everolimus, temsirolimus, and bortezomib at their respective IC50s after 48 h of treatment. The latter drug is a proteasome inhibitor and was used as a positive control. Hierarchical clustering analysis of the data revealed significant differences between the molecular fingerprints of rapamycin and its analogs. Specifically, the cellular response to rapamycin and temsirolimus were similar, but that of everolimus differed dramatically from both of them (Figure 1, A). On the other hand, the effects of everolimus and bortezomib were alike, suggesting the possible involvement of everolimus in inhibition of the proteasome. Furthermore, 740 proteins, majority of which are specifically up-regulated in the samples treated with everolimus and bortezomib (cluster 8 in Figure 1, A), mapped to proteasome-related GO terms in the enrichment analysis (Figure 1, B and C).

**Figure 1.**
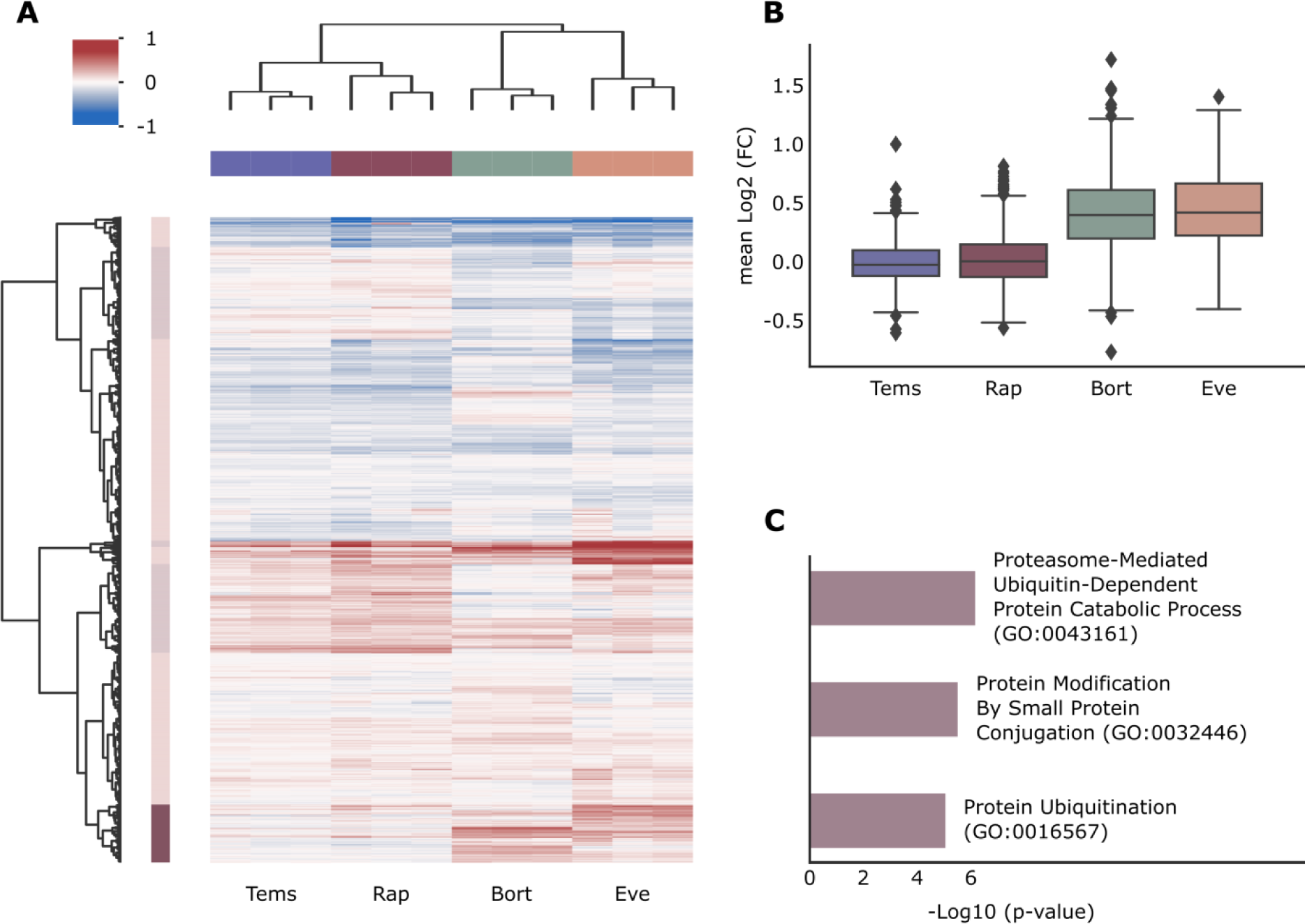
(A) Hierarchical clustering of proteomics data of A549 cell line treated with four drugs showing similarity between proteome signatures of everolimus- and bortezomib-treated samples. Proteins clustered into eight main groups; cluster 8 is highlighted in dark red. (B) Distributions of mean protein fold changes of the protein cluster 8 and (C) major enriched GO terms of the protein cluster 8.

To evaluate the effect of each drug on the cell line proteome, statistical t-testing with Benjamini-Hochberg (BH) multiple testing correction was employed, and volcano plots were generated for all four drugs. The number of outliers was determined using the threshold values of fold change <0.5 or >2 and FDR BH <0.05, as depicted in Figure 2. The drug with the fewest outliers is temsirolimus, followed by rapamycin, while bortezomib and particularly everolimus produced most outliers, suggesting that the latter two drugs had the highest impact on the proteome. All test results are summarized in Supplementary Table S2.

**Figure 2.**
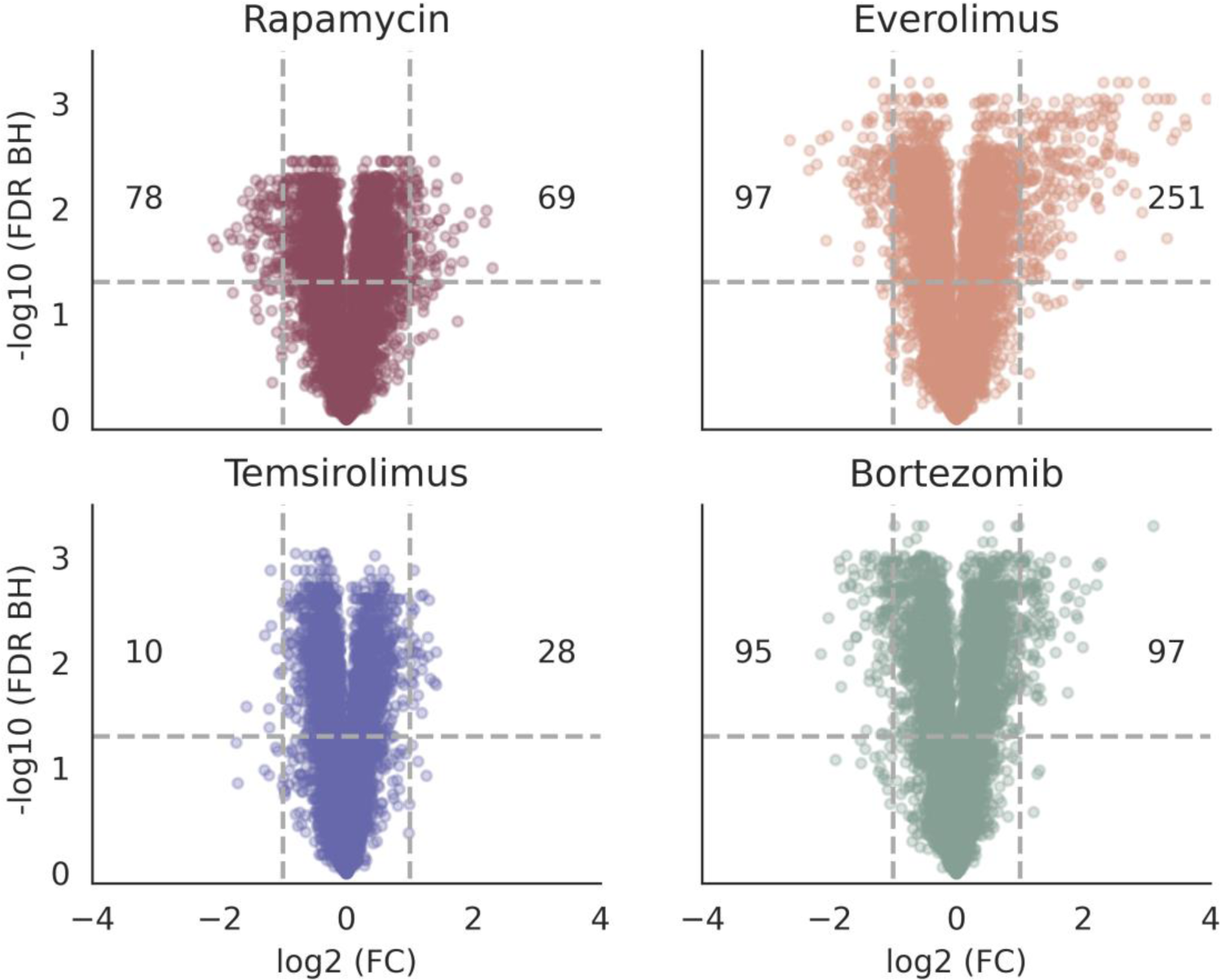
Volcano plots showing the disturbance of A549 cell line proteome by four studied drugs. The numbers of outliers are shown for the following thresholds: fold change > 2 or <0.5, FDR BH < 0.05.

Figure 3 shows the numbers of shared outliers between different drugs, upregulated proteins are shown with the shades of red, while the down-regulated ones are depicted with the shades of blue. The Venn diagrams showing the shared outliers are given in Supplementary Figure S1. The largest numbers of both positive and negative outliers are obtained for everolimus and bortezomib, consistent with these two drugs having similar action mechanisms. It is worth noting that in terms of protein outliers, rapamycin and bortezomib treatments share much more similarity than was observed in the hierarchical clustering of the whole proteome.

**Figure 3.**
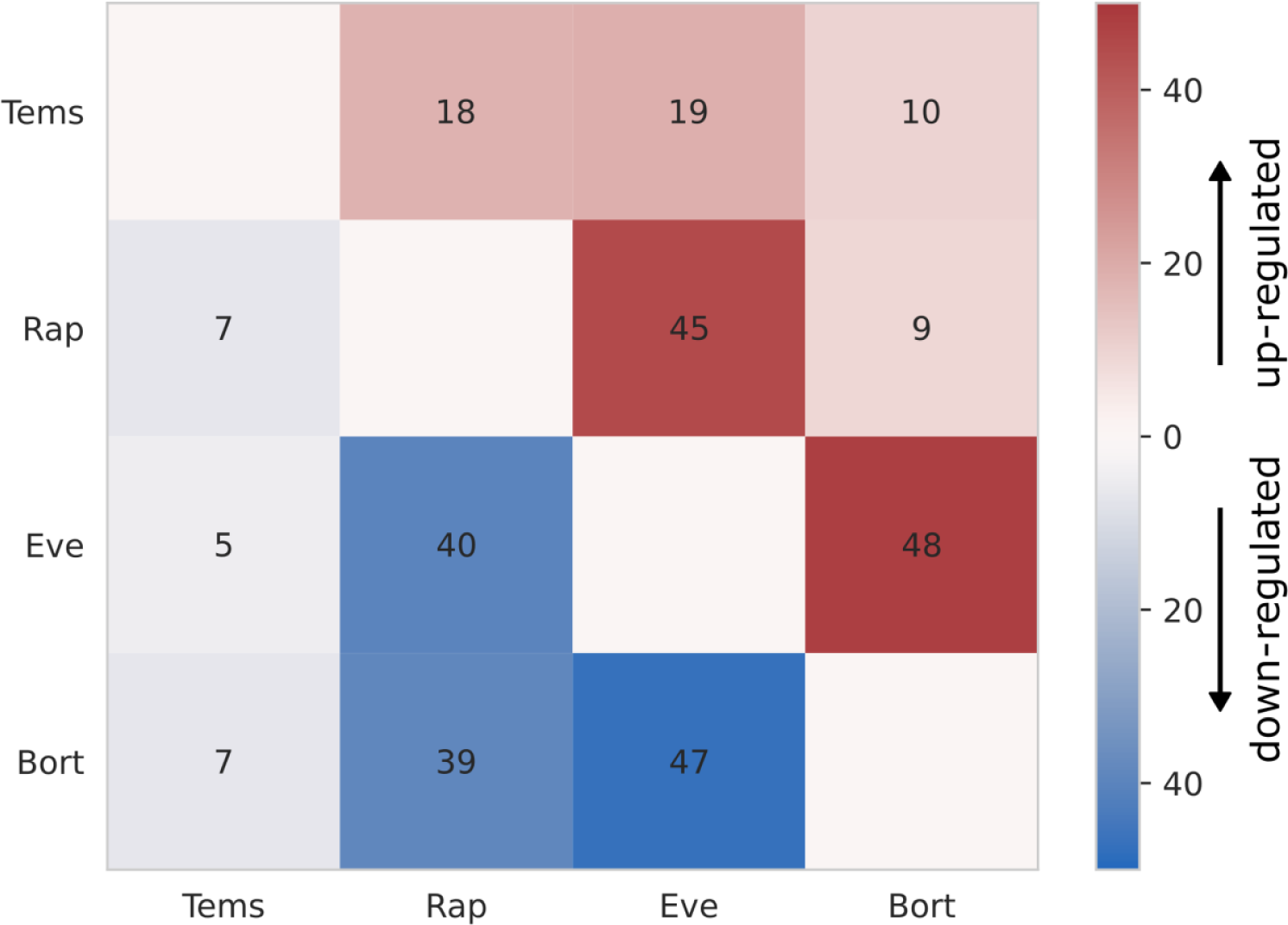
Number of shared protein outliers between different treatments. Up-regulated outliers are shown as shades of red, and down-regulated ones with shades of blue.

GO enrichment was performed for the outlier lists for each of the drugs, and the results are summarized in Supplementary Table S3. The terms related to UPS were enriched in the outlier gene sets corresponding to bortezomib and rapamycin treatment, but not for everolimus. The analysis of the shared GO terms (Supplementary Figure S2 A) revealed significant similarity between the GO terms affected by rapamycin and bortezomib treatments. The sets of up- and down-regulated proteins for these two drugs were then used separately for the GO enrichment, and the comparison of the resulting terms (Supplementary Figure S2 B) has shown that the down-regulated proteins are mostly responsible for the similarity. The corresponding GO terms are listed in Supplementary Table S4. Among the shared GO terms, the following are related to UPS: Positive Regulation of Ubiquitin Protein Ligase Activity (GO:1904668), Positive Regulation of Ubiquitin-Protein Transferase Activity (GO:0051443), and Regulation of Ubiquitin Protein Ligase Activity (GO:1904666). Note, however, that GO-term enrichment analysis of the outliers did not lead to similarly enriched pathways, perhaps indicating that the identity of the proteins affected by everolimus is distinct from that of bortezomib.

### Polyubiquitination patterns

Protein polyubiquitination is a primary and crucial stage of proteasome-mediated protein degradation. Ubiquitin molecules can form polyubiquitin chains by covalently binding to each other in one of eight sites: N-terminus, K6, K11, K27, K29, K33, K48, and K63.^63,64^ Among these sites, the K48 linkage is the most frequently associated with proteasomal degradation, although there is evidence that all other lysine-linked polyubiquitin chains may also serve as degradation signals for proteasome.^65,66^

In order to assess the effect of the drugs on the UPS, we searched the proteomic data for the polyubiquitination patterns by performing the search over a limited sequence database. The limited database included a set of ubiquitin sequences with GG added to all potential ubiquitination sites (Supplementary Table S1). Trypsin cleavage of ubiquitin-modified protein results in diglycine (GG) modification of the corresponding lysine residue (K), which cannot be directly modified by the TMT: the latter binds to the N-terminus of GG instead. The use of a limited search space increases the search sensitivity at the price of a higher probability of false matches. To reduce the latter, all search results were filtered to exclude C-terminal position of GG-modified K residues, since trypsin cannot cleave the peptide bond after the modified residue.

For each potential modification site, both modified and unmodified peptides were quantified. Peptides not containing any potential ubiquitination site were quantified separately. Figure 4 shows the results of this analysis: the overall ubiquitin abundance was significantly increased for both bortezomib- and everolimus-treated samples, while rapamycin and temsirolimus treatment resulted in only a marginal increase (Figure 4). The GG-modified peptides also demonstrated similar polyubiquitination patterns for both bortezomib and everolimus, specifically, exhibiting high abundance of K11 and K48 linkage associated with proteasomal protein degradation. K63-linked polyubiquitin appears more pronounced for everolimus treatment, while the K29-linked one is more abundant in the bortezomib-treated sample. For the two remaining compounds, the K48 linkage was also observed, even thoughthe ubiquitin abundance up-shift was barely present.

**Figure 4.**
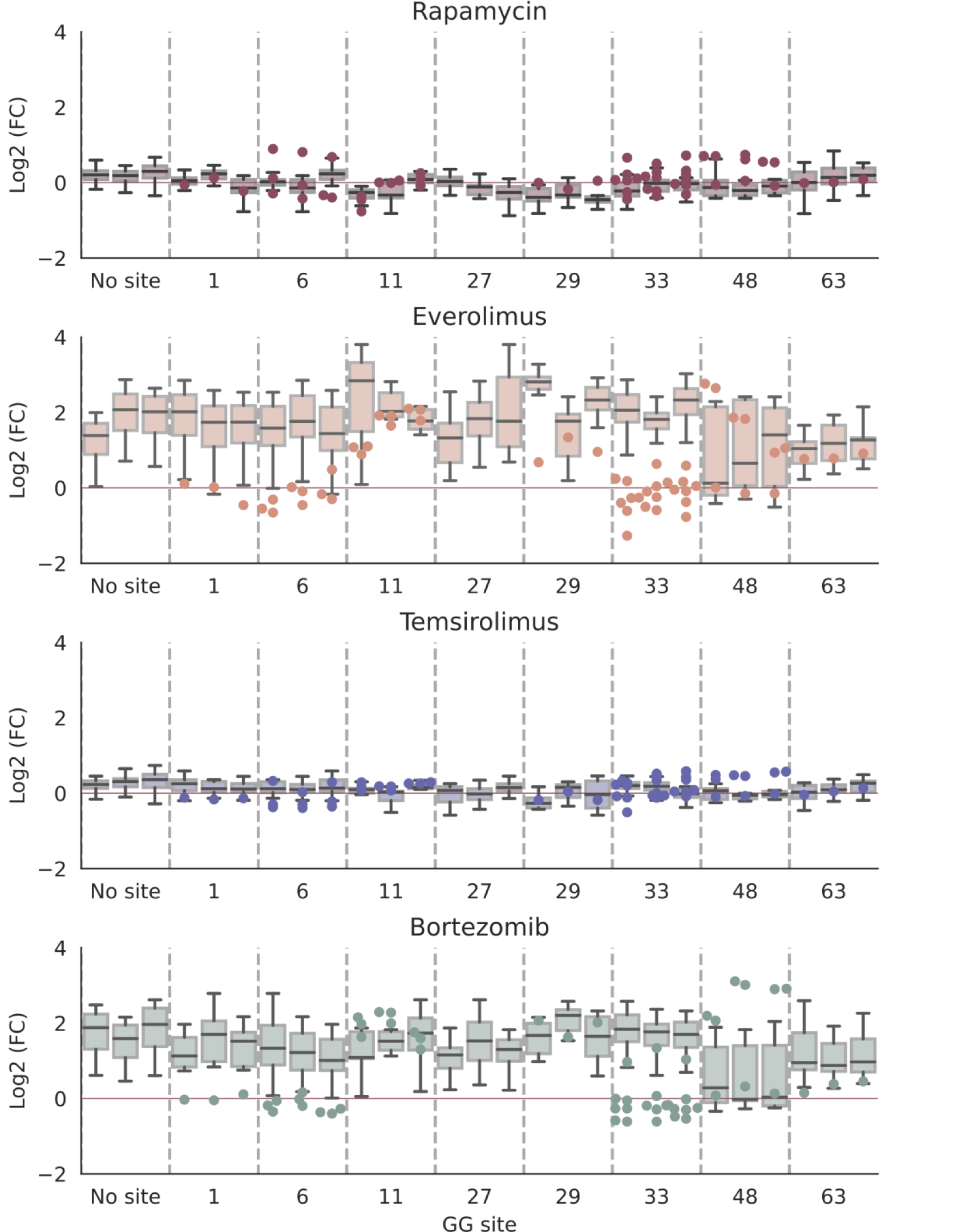
Fold changes of ubiquitin peptides obtained from the limited database search (see the text). The peptides were grouped based on the appearance of GG-modification sites. FCs of unmodified peptides are shown as box plots, while those of the GG-modified ones are presented as dots, each dot representing one peptide-spectrum match (PSM).

A recently discovered mechanism of regulation of UPS is via enzymatic phosphorylation of ubiquitin by PINK1 at residue S65 ^67–69^. This modifications results in changes in the conformational states of polyubiquitin, further impacting the quaternary arrangements of polyubiquitin subunits and, thus, inhibiting the activities of enzymes responsible for attaching and removing polyubiquitins^70,71^. This mechanism is also known as a mitophagy initiation signaling^72^. The limited ubiquitin search of the proteomic data under study revealed differentially abundant S65 phosphorylation of ubiquitin (Figure 5). Specifically, we found that the everolimus treatment resulted in significant elevation of ubiquitin phosphorylation, indicating a unique interplay between phosphorylation signaling and UPS triggered by this drug.

**Figure 5.**
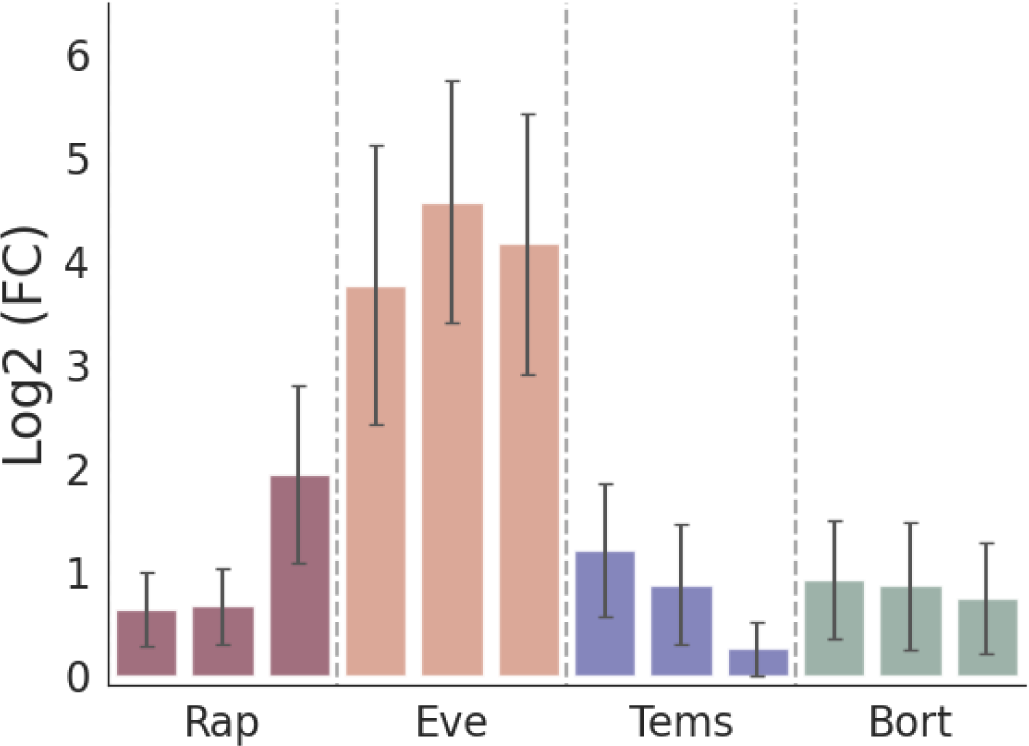
Fold changes of S65-phosphorylated ubiquitin. Bars show mean value of the three corresponding PSMs, while the error bars show the standard deviation.

The stoichiometry of ubiquitin phosphorylation in the control sample was estimated using the following formula:

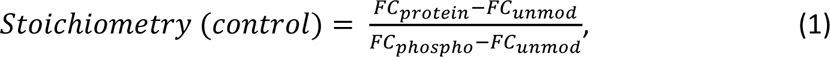

where FC stands for fold changes between control and everolimus-treated samples of the protein (*FC*_*protein*_), S65-containing unmodified (*FC*_*unmod*_) and phosphorylated peptide (*FC*_*phospho*_). Protein fold change was estimated as median FC of all ubiquitin peptides without any GG- or phospho-modification site. Stoichiometry upon drug treatment is then calculated as follows:

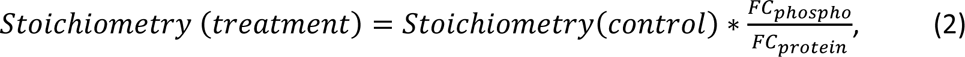

where the fold changes correspond to the treatment in question. The stoichiometries of S65 phosphorylation were 13, 21, 45, 15, and 5% in the control sample and after treatment with rapamycin, everolimus, temsirolimus, and bortezomib, respectively. Everolimus treatment led to the highest phosphorylation rate, illustrating specific cellular mechanism invoked by this drug.

### Polyubiquitination in the ProTargetMiner dataset

For additional verification of our findings, the same limited ubiquitin search approach was then applied to an earlier published ProTargetMiner^58^ dataset, including expression proteomic analysis of A549 cell line treated by 56 anticancer compounds. Supplementary Figure S3 shows the summary of the results obtained for all experiments. The threshold value for statistically significant fold change (compared to untreated controls) of ubiquitin level set at 2 was exceeded for four drugs: auranofin, bortezomib, b-AP15, and everolimus. The results for the first three drugs were expected as they are known as proteasome inhibitors.^73–77^ Everolimus treatment resulted in a significantly elevated ubiquitin level, further supporting the results of this study. Also, temsirolimus included in the panel of drugs analyzed in ProTargetMiner study did not change the level of ubiquitin in the cells.

The results of the search using the generated GG-peptide database are shown in Supplementary Figure S4 as relative intensities (log2 fold change compared to untreated control) of peptide-spectrum matches (PSMs) corresponding to GG-modified and phosphorylated ubiquitin peptides across the panel of 56 drugs. Since the multi-batch design of ProTargetMiner dataset was prone to missing values, only K29 and K48 linkages were universally identified for most drug treatments considered and were included in the plot for the sake of clarity. In general, these modifications correlate with the total amount of ubiquitin in samples (shown as horizontal lines), with a small number of outliers probably corresponding to false matches. A significant up-regulation of K48-linked ubiquitin after everolimus treatment indicates that a significant portion of the excessive ubiquitin corresponds to polyubiquitination of the proteins tagged for proteasomal degradation. Supplementary figure S4 also shows that the concentration of S65-phosphorylated ubiquitin is clearly elevated in response to certain drugs, including everolimus further supporting the results of this study. An extremely elevated phosphorylated ubiquitin was also observed for the sorafenib treatment, which is a known protein kinase inhibitor.^78^ At the same time, the ubiquitin phosphorylation level for sorafenib treatment does not correlate with the elevated total ubiquitin concentration or K48-linked polyubiquitin. Regarding the rapalogs, the observations from ProTargetMiner dataset seem very similar to our main dataset.

### Proteolytic activity observed in the everolimus-treated sample

Since UPS system is responsible for protein degradation, it is possible to detect the degradation products in the proteomes by performing a database search with semi-tryptic protease specificity. The identified semi-tryptic peptides are of particular interest if they have significantly high fold changes (FC) while the fold changes of corresponding proteins are low. Figure 6, A shows the FC of the identified semi-tryptic peptides and the corresponding proteins in the everolimus-treated samples. A distinguished group of up-shifted semi-tryptic peptides was observed exclusively for this drug treatment, suggesting that the treatment induced some protease activity, which may or may not be attributed to the proteasome. The group of 174 up-shifted semi-tryptic peptides marked by red dots in Figure 6, A was selected using the following filters: FC >2 for semi-tryptic peptide, FDR BH< 0.05, FC < 2 for the corresponding protein. Plots for all four drugs are given in Supplementary Figure S5.

**Figure 6.**
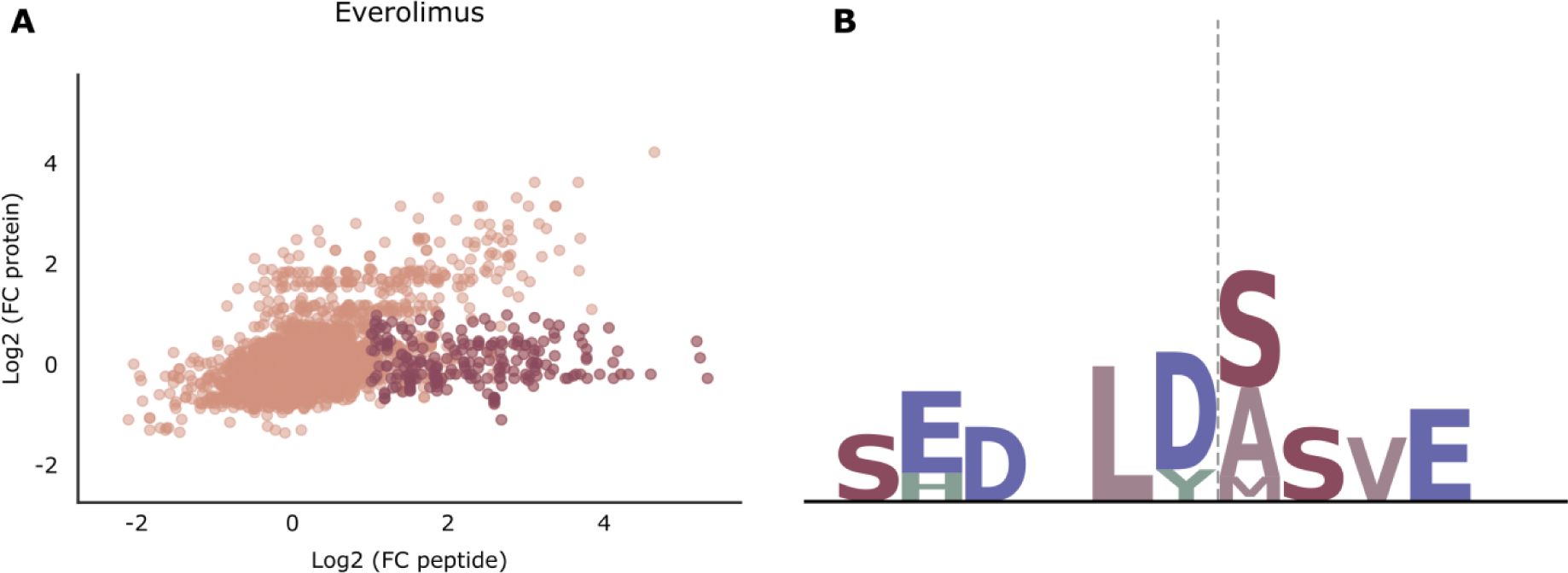
(A) Fold changes (log2-transformed) of semi-tryptic peptides identified in the sample treated with everolimus and the corresponding proteins. A group of 174 peptides depicted as red dots was selected using the following filters: FC >2 for semi-tryptic peptide, FDR BH < 0.05, FC < 2 for the corresponding protein. (B) Motif enrichment performed for the set of semi-tryptic peptides up-shifted upon everolimus treatment. Vertical dashed line shows the cleavage site. The DXLD motif similar to that of several caspases^71^ is observed.

We performed motif enrichment for this set of 174 peptides, corresponding to 167 unique cleavage sites and found 32 of them corresponding to a cleavage at aspartate residue. Moreover, the clear DXLD motif N-terminal to the cleavage site was enriched for this set of semi-tryptic peptides (Figure 6, B), which is similar to the cleavage specificity of some caspases ^79^. GO enrichment of the corresponding gene set revealed mostly structural proteins (Supplementary Table S5). Since the nature of this proteolytic activity is unknown, one possible explanation is that it is due to the proteasomal degradation. In order to test this assumption, another limited search for GG-modified peptides was performed on a subset of the proteins corresponding to the above 174 peptides. The output was validated manually, and GG-modification was found for Heat shock cognate 71 kDa protein (HSCP7C). Despite the fact that this protein is highly ubiquitinated in both everolimus- and bortezomib-treated samples (Supplementary Figure S6, A), only everolimus treatment resulted in significant up-shift of its semi-tryptic peptides (Supplementary Figure S6,B). This indicates that everolimus may indeed invoke the proteasomal degradation of this protein, while bortezomib inhibited the proteasome leaving HSP7C ubiquitinated and not cleaved.

Proteolytic activity invoked by everolimus treatment may be associated with both caspases, or other proteases, and the proteasome. It is indicative of the unique mechanism of action of this drug that is strikingly different from that of rapamycin, temsirolimus, and even of bortezomib, with which it otherwise shares many similar features.

Note that the differences in treatment outcome between everolimus and temsirolimus revealed in this proteomic study have been noted in clinical practice before, indicating indirectly the differences in their pharmacodynamics. Namely, everolimus treatment of metastatic renal cell carcinomapatients resulted in higher overall survival as compared to temsirolimus treatment. This difference is either due to the different mechanisms of action as our results demonstrated, or the drug administration guidelines.^7,8^ Indeed, these two drugs are prescribed following different indications: everolimus is recommended for patients previously treated with an anti-VEGFR tyrosine kinase inhibitor, while temsirolimus is recommended as first-line treatment in patients with poor-risk features. Also, there exists an assumption about the interconnection of mTOR and NFkB pathways including proteasome activity regulation ^80^ that may be responsible for the observed effects. A synergistic therapeutic effect was also shown for a combination of everolimus with bortezomib in multiple myeloma cells where inhibition of the AKT/mTOR pathway was presented as the potential mechanism.^81^

## V. Conclusion

The mechanisms of action of rapamycin and its analogs, everolimus and temsirolimus, were investigated using expression proteomics, with a particular focus on the UPS activity. Interestingly, the proteomes affected by everolimus and bortezomib showed a striking similarity, especially in the expression changes of proteins related to UPS pathway. The level of ubiquitin and its tryptic peptides with GG modification of K11 and K48 residues indicated similar polyubiquitination patterns resulting from these two treatments. However, no significant changes in ubiquitin abundance were observed for rapamycin and temsirolimus in either the ProTargetMiner data or the separate experiment performed in this study. Despite this, GO enrichment analysis did reveal a similarity between the sets of genes down-regulated by both rapamycin and bortezomib, including some UPS-related terms, perhaps indicating that a unique population of proteins is affected by each drug. Another interesting observation was the significantly elevated S65-phosphorylation of ubiquitin in case of evelolimus, which may further shed light on the mechanism of its action. This finding is similar to the one observed for sorafenib, which inhibits multiple intracellular serine/threonine kinases. Additionally, everolimus treatment resulted in increased proteolytic activity, which could be associated with both the proteases and the proteasome, providing further evidence of its distinct mechanism of action. However, no specific up-regulation upon everolimus treatment was observed for any of the caspases. Overall, proteomic data suggest the involvement of UPS in the mechanism of action of everolimus and stresses the difference between the actions of chemically analogous compounds, highlighting the need for further studies to better understand the pharmacodynamics of rapalogs. Overall, these findings might help explain the distinct clinical indications of these drugs in disease therapy.

## Acknowledgments

This work was supported by Russian Science Foundation (continuation project #20-14-00229 to M.V.G.). R.A.Z. also acknowledges support from The Ministry of Science and Higher Education of the Russian Federation, agreement no. 075-15-2020-899. The authors have declared no conflict of interest.

## Notes

### Competing Interest Statement

The authors have declared no competing interest.

