## Supplementary Figure S1 for "Chemical proteomics reveals that the anticancer drug everolimus affects ubiquitin-proteasome system"

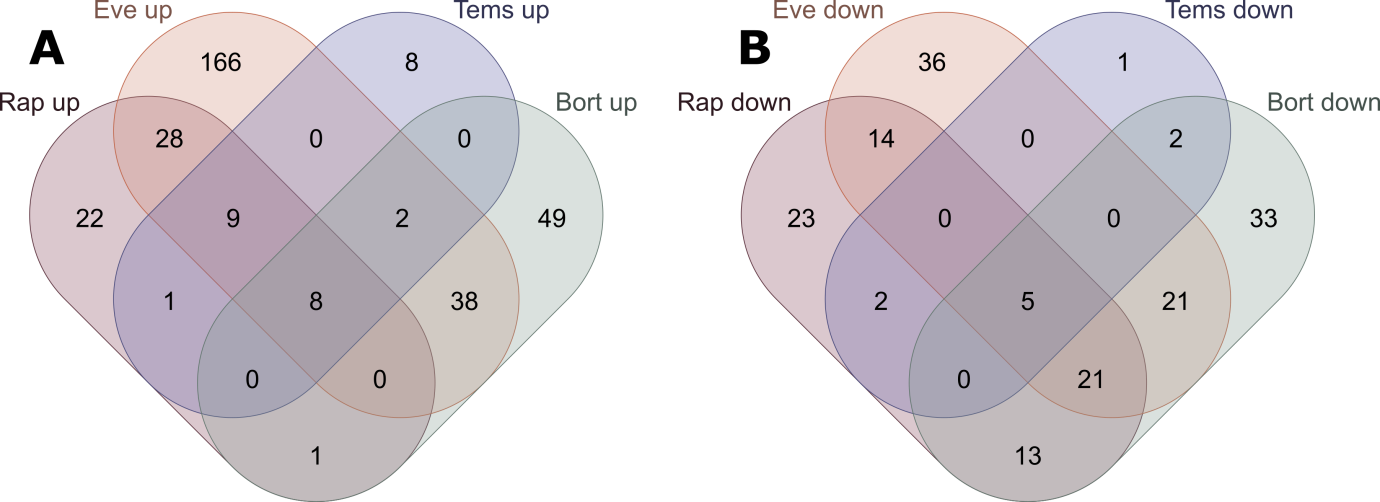

**Supplementary figure S1.** Number of shared up-regulated and down-regulated protein outliers between different treatments. No up-regulated outlier from one treatment was down-regulated in a different treatment.
