## Supplementary Figure S2 for "Chemical proteomics reveals that the anticancer drug everolimus affects ubiquitin-proteasome system"

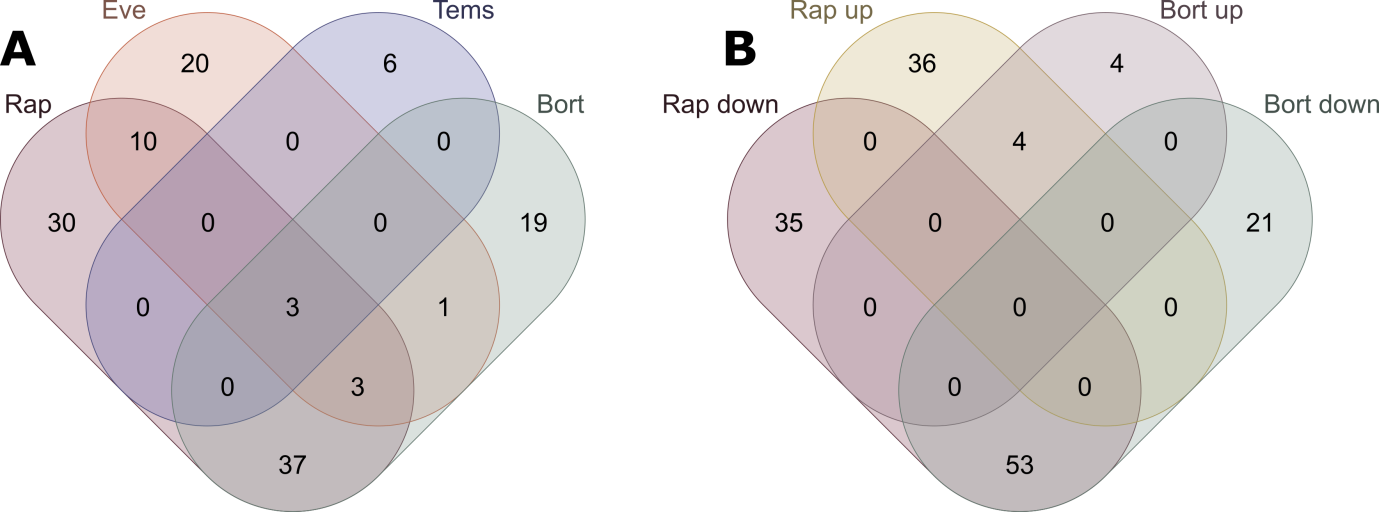

**Supplementary Figure S2.** (A) Number of shared GO terms enriched based on up- and down-regulated outliers in the proteomes of A549 cells treated with four drugs. (B) Number of shared GO terms enriched separately for up- and down-regulated proteins in the samples treated with rapamycin and bortezomib.
