## Supplementary Figure S3 for "Chemical proteomics reveals that the anticancer drug everolimus affects ubiquitin-proteasome system"

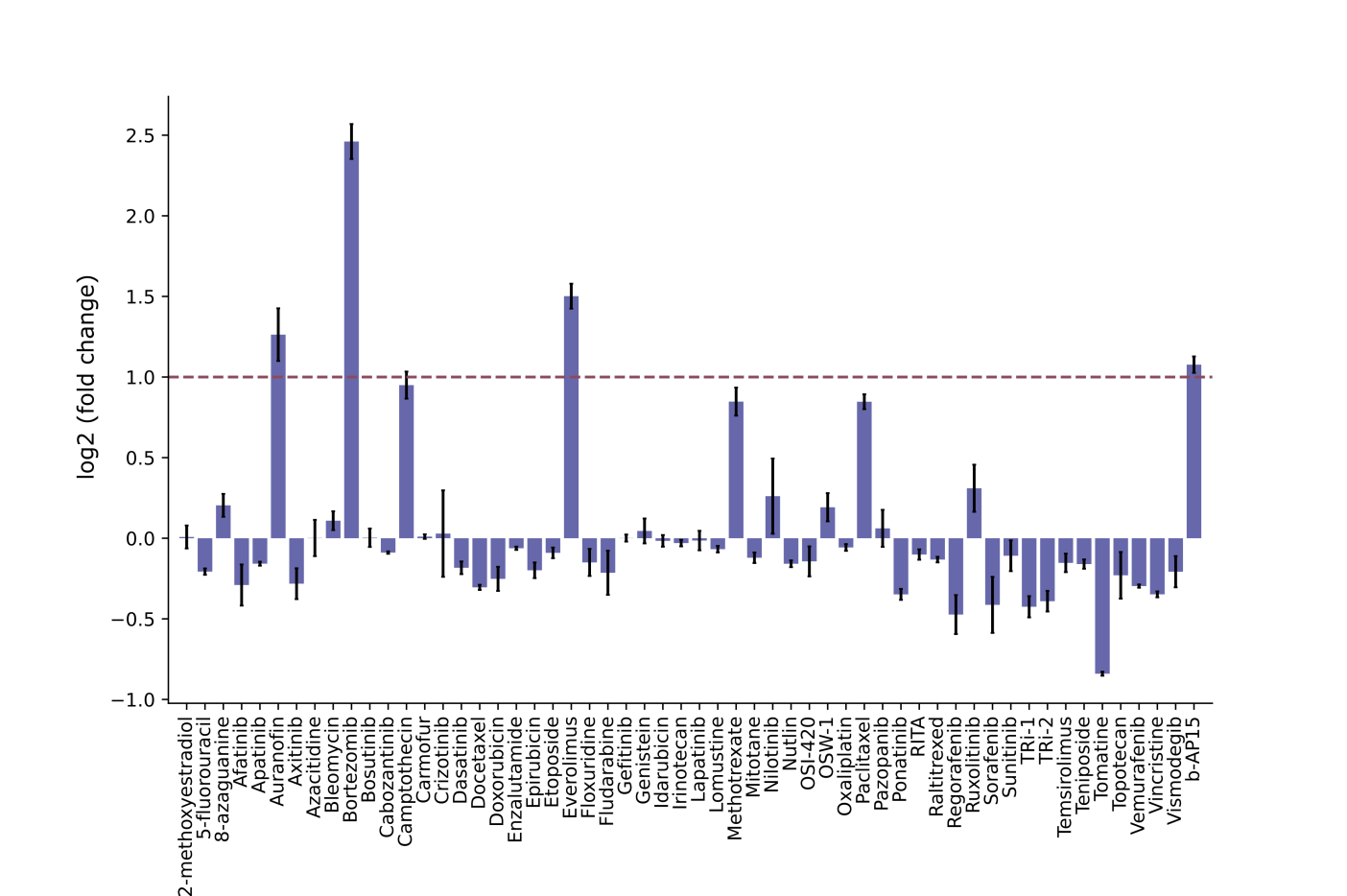

**Supplementary Figure S3.** The relative quantity of ubiquitin (as log2–transformed fold change relative to control) for A549 cell line treated with 56 anticancer compounds from ProTargetMiner dataset.
