## Supplementary Figure S4 for "Chemical proteomics reveals that the anticancer drug everolimus affects ubiquitin-proteasome system"

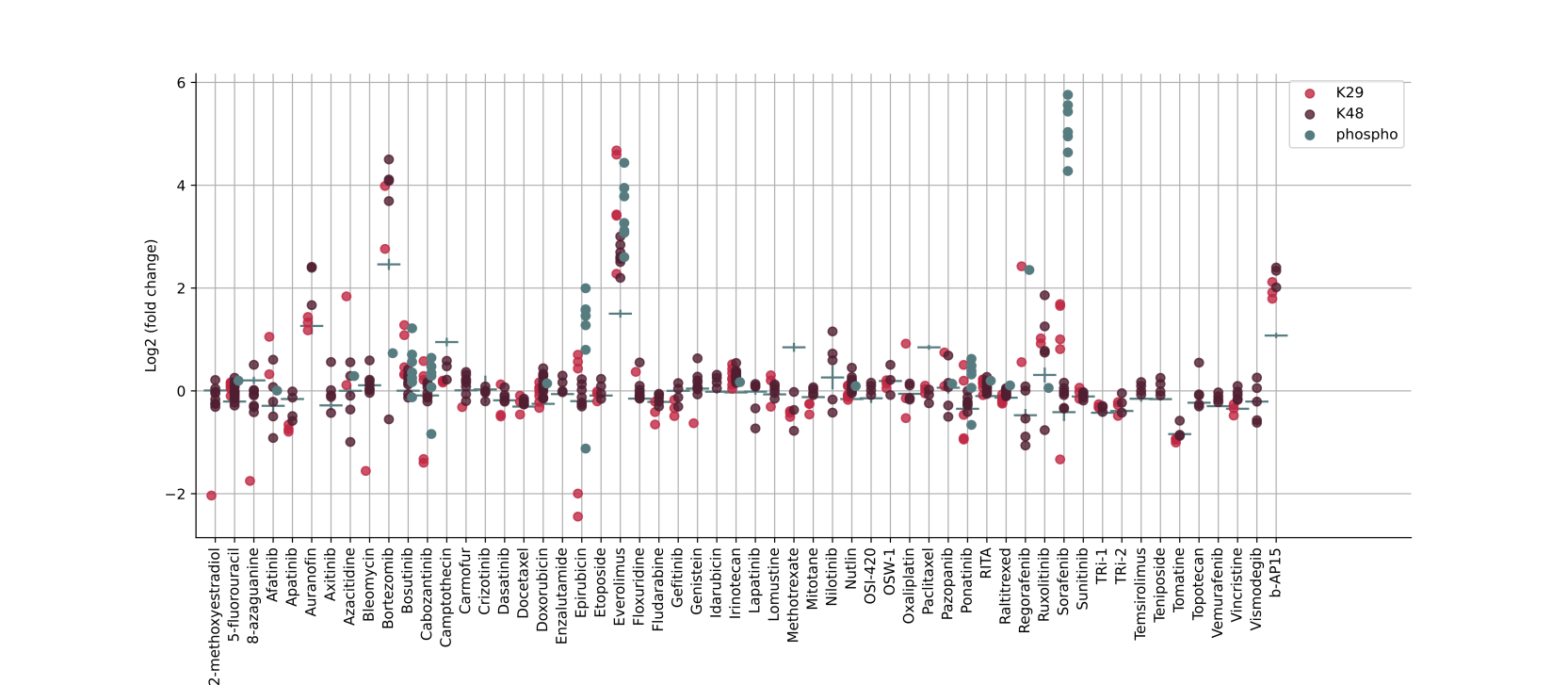

**Supplementary Figure S4.** Relative intensities of PSMs corresponding to ubiquitinated and phosphorylated ubiquitin in the samples treated with 56 chemotherapeutic drugs.
