## Supplementary Figure S5 for "Chemical proteomics reveals that the anticancer drug everolimus affects ubiquitin-proteasome system"

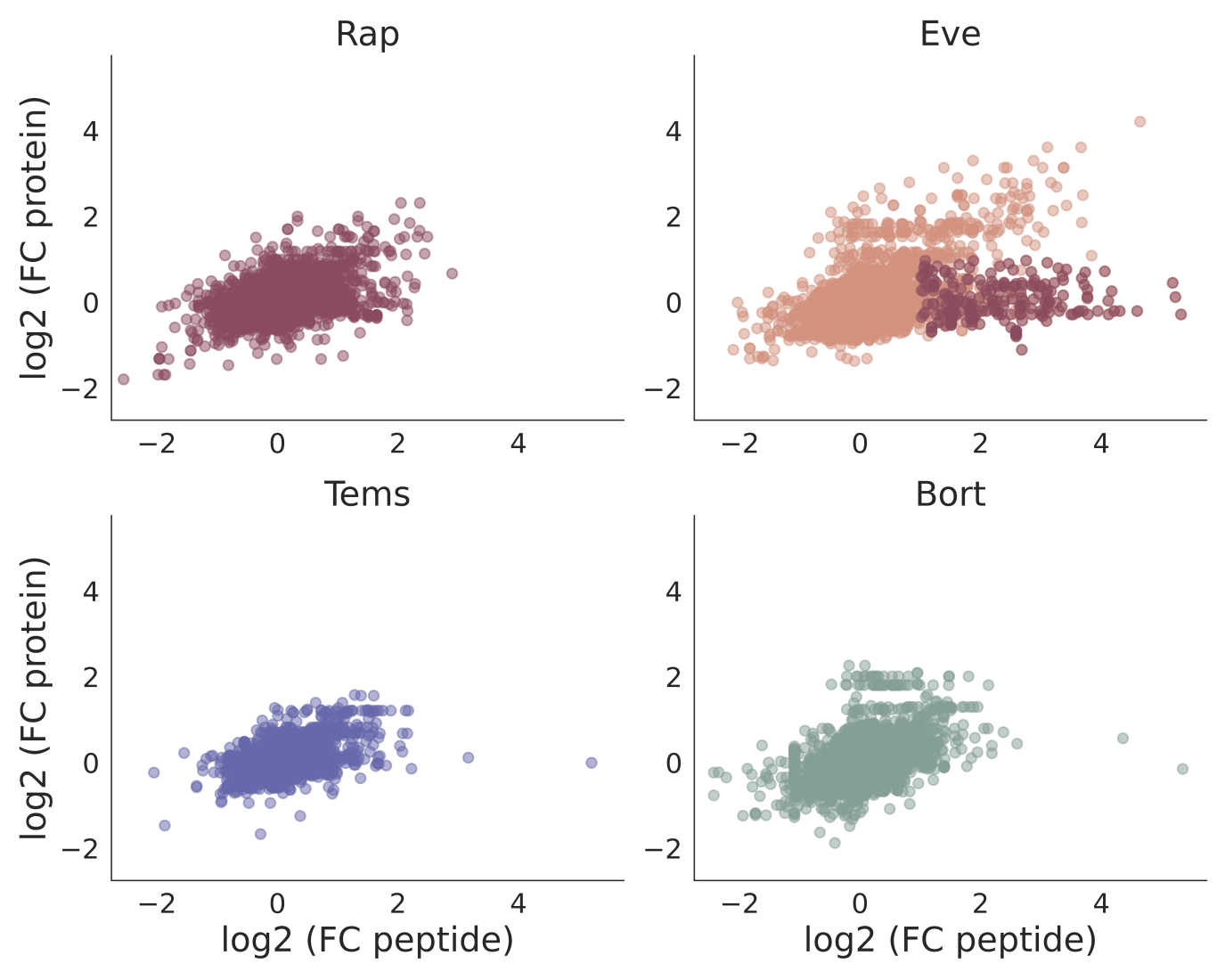
**Supplementary Figure S5.** Fold changes (log2-transformed) of semi-tryptic peptides identified in the sample treated with four drugs and the corresponding proteins. A significant group of up-shifted semi-tryptic peptides was observed only for the everolimus treatment (marked as red dots), indicating the induction of protease activity by this drug.
