## Supplementary Figure S6 for "Chemical proteomics reveals that the anticancer drug everolimus affects ubiquitin-proteasome system"

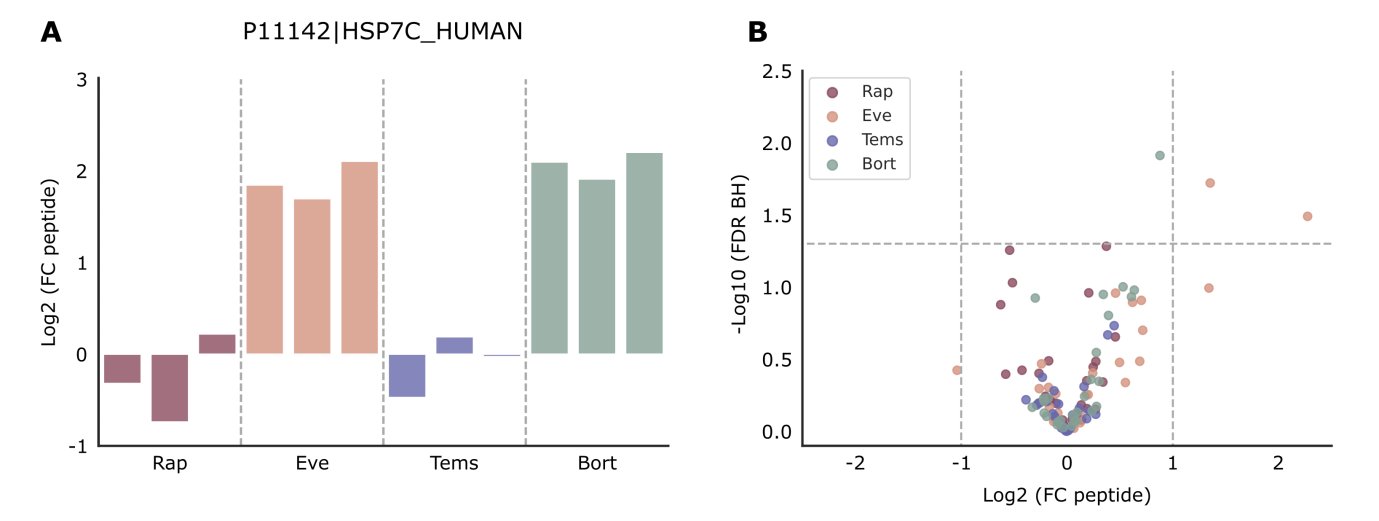

**Supplementary Figure S6.** (A) Fold changes of GG-modified peptide SINPDEAVAYGAAVQAAILSGD***K***SENVQDLLLLDVTPLSLGIETAGGVMTVLIKR from HSP7C protein (modified K residue shown in bold italic), (B) volcano plot of semi-tryptic peptides corresponding to this protein. Significantly up-shifted peptides observed only for the everolimus treatment.
